# Device-free isolation of photoreceptor cells from patient iPSC-derived retinal organoids

**DOI:** 10.1101/2024.05.02.592255

**Authors:** Nicholas E. Stone, Laura R. Bohrer, Nathaniel K. Mullin, Jessica Cooke, Allison T. Wright, Edwin M. Stone, Robert F. Mullins, Budd A. Tucker

## Abstract

Autologous photoreceptor cell replacement therapy shows great promise for treating patients with multiple forms of inherited retinal degenerative blindness. Specifically, in disorders such as retinitis pigmentosa and Stargardt’s disease, selective death of photoreceptor cells results in irreversible blindness. Induced pluripotent stem cell (iPSC) derived retinal organoids, which faithfully recapitulate the structure of the neural retina, are an ideal source of photoreceptor cells required for these therapies. However, in addition to photoreceptor cells, retinal organoids also contain many other retinal cell types. Therefore, approaches for isolating fate committed photoreceptors from dissociated retinal organoids are desirable to produce photoreceptor cell replacement therapies. In this work, we present a partial dissociation strategy, which leverages the high level of organization found in retinal organoids to enable selective enrichment of photoreceptor cells without the use of specialized equipment or reagents such as antibody labels. Given that this technique can be performed with only standard plasticware and cGMP compliant reagents, it is an ideal candidate for use in the preparation of clinical cell therapies.

## Introduction

In several forms of inherited retinal degeneration, gradual loss of photoreceptor cells in a patient’s retina leads to progressive vision loss. While gene therapy shows promise for halting disease progression provided a molecular diagnosis is available, restoration of vision will likely require photoreceptor cell replacement (1–8). One promising source of transplantable photoreceptors are induced pluripotent stem cell (iPSC)-derived retinal organoids. We and others have developed 3D retinal differentiation protocols which produce retinal organoids that recapitulate the structure of the human retina and contain fate-committed photoreceptor cells (9–19). Our group is particularly interested in using patient-derived iPSC lines to produce autologous photoreceptor therapies, which we believe will eliminate the need for long-term immune suppression following transplantation. We have produced current good manufacturing practice (cGMP) protocols for both the generation and differentiation of patient-derived iPSCs and are exploring methods by which production can be scaled to clinically-relevant levels in an academic environment (9, 10, 20).

While clinical-grade production of patient-derived retinal organoids containing transplantable photoreceptor precursor cells is now possible, to enhance potency of the therapeutic product a method for isolating photoreceptor cells is required. Specifically, transplantation of inner retinal neurons, which remain in many patients with inherited retinal degeneration, is not likely to contribute to the desired clinical outcome and may interfere with photoreceptor integration with the remaining neurons. Fluorescence activated cell sorting (FACS) and magnetic activated cell sorting (MACS) constitute gold standard cell sorting technologies. Both are mature, high throughput and exhibit very high selectivity provided antibody labels can be targeted to unique cell surface antigens (or a combination of markers in the case of FACS) present on cells selected for enrichment or depletion (21–25). However, using animal-derived antibodies to target fluorescent probes or magnetic beads is not an ideal approach when isolating cells destined for human transplantation. Further, processing multiple patient samples on single pieces of large, expensive equipment increases the risk of cross contamination as well as the cost of the resulting therapy. Thus, other, GMP-compliant methods for enriching photoreceptor cells from non-photoreceptors would be highly desirable.

Retinal organoids faithfully reproduce the architecture of the mature human retina and exhibit a characteristic laminar structure in which the photoreceptors are found in the outermost layer. In this work, we evaluated the degree to which partial dissociation can be used to exploit this spatial segregation to enrich transplantable photoreceptor cells from patient iPSC-derived retinal organoids. Such an approach would decrease the regulatory burden of the resulting cell therapy by removing the need for xenobiotic sorting reagents, reduce cost by avoiding the use of specialized equipment and increase the potency of the resulting cell therapy by enriching the cells which contribute to the desired clinical outcome.

## Methods

### Patient-derived iPSC generation and validation

This study was approved by the Institutional Review Board of the University of Iowa (project approval #200202022) and adhered to the tenets set forth in the Declaration of Helsinki. Three patient lines were used in this study. Patient iPSCs were generated from an individual with no disease (Line 2: B1427) and 2 individuals with molecularly confirmed enhanced s-cone syndrome (Line 1: B342cor (homozygous for c.119-2A>C mutation in *NR2E3*) and Line 3: B1737 (compound heterozygous for c.932G>A; p.(Arg311Gln) and c.119-2A>C mutations in *NR2E3*) (9, 26). For Line 1, the disease-causing mutation in NR2E3, c.119-2A>C, was corrected via CRISPR-mediated homology dependent repair in patient-derived iPSCs as described previously (26). Line 3 was sorted into well-organized (laminated neural retina with few appendages) and poorly-organized (poor lamination, excessive aggregation, or presence of none-retinal tissue) populations before performing partial dissociation.

### Retinal organoid differentiation

Retinal differentiation was performed as described previously with minor modifications (26, 27). Briefly, iPSCs were cultured on laminin 521 coated plates in E8 medium. Embryoid bodies (EBs) were lifted with ReLeSR (STEMCELL Technologies, Cambridge, MA) and transitioned from E8 to neural induction medium (NIM) over a four-day period. On day 6, NIM was supplemented with 1.5 nM BMP4 (R&D Systems, Minneapolis, MN). On day 7, EBs were adhered to CELLstart coated plates (Thermo Fisher Scientific). BMP4 was gradually transitioned out of the NIM over seven days. On day 16, the media was changed to retinal differentiation medium (RDM). On day 25-30 the entire EB outgrowth was mechanically lifted and transferred to ultra-low attachment flasks in 3D-RDM (RDM plus 10% KnockOut Serum Replacement (KSR)); Thermo Fisher Scientific), 100 μM taurine (Sigma-Aldrich), 1:1000 chemically defined lipid concentrate (Thermo Fisher Scientific), and 1 μM all-trans retinoic acid (until day 100; Sigma-Aldrich). The cells were fed three times per week with 3D-RDM until harvest at Day 160.

### Staged partial dissociation of retinal organoids

For each of the 3 samples tested, approximately 20 organoids were selected for staged partial dissociation along with 10 organoids which were dissociated in one step as a control. Both populations were placed in 1.5mL Eppendorf tubes and suspended in 0.5mL of 30U/mL papain (Worthington) and 120U/mL DNase (Worthington) in Earle’s balanced salt solution (EBSS) (Worthington). These tubes were then placed in a rotating incubator at 37°C. At 20, 40 and 60 minutes, the organoids being partially dissociated were allowed to settle, after which the supernatant containing liberated cells was collected. These organoids were washed once with papain and then placed in fresh papain solution for continued dissociation. The cells in the supernatant were passed through a 100 μm filter and counted using a Moxi Go II coulter counter (Orflo). After 60 minutes of total dissociation, any remaining solid aggregates in both the experimental and control tubes were fully dissociated via trituration (i.e., mechanical aspiration and expulsion via a 1mL pipette), counted, pelleted and prepared for scRNA-seq. All samples were pelleted and resuspended in 8μg/mL recombinant albumin (New England Biolabs) in dPBS[-/-] for encapsulation with the Chromium X instrument (10X Genomics). Approximately 8,000 cells were targeted for encapsulation per sample.

### Single-cell gene expression library preparation and sequencing

Single cells were partitioned and barcoded with the Chromium Controller instrument (10X Genomics) and Single Cell 3’ Reagent (v3.1 chemistry) kit (10X Genomics) according to the manufacturer’s specifications with no modification (Rev E). Final libraries were quantified using the Qubit dsDNA HS Assay Kit (Life Technologies) and diluted to 3ng/μL in buffer EB (Qiagen). Library quality was confirmed using the Bioanalyzer High Sensitivity DNA Assay (Agilent) prior to sequencing as previously described (28).

### Single-cell gene expression data integration and processing

scRNA libraries were pooled and sequenced using the NovaSeq 6000 instrument (Illumina) generating 100-bp paired end reads. Sequencing was performed by the Genomics Division of the Iowa Institute of Human Genetics. FASTQ files were generated from base calls with the bcl2fastq software (Illumina), and reads were mapped to the pre-built GRCh38 reference (refdata-gex-GRCh38-2020-A) with Cell Ranger v7.0.0 (10X Genomics) using the ‘count’ function with the following parameters: --expect-cells=8000 --localcores=56. Only cells passing the default Cell Ranger call were analyzed further. All timepoints and control dissociations for the four samples studied (Line 1, Line 2 and the well-organized and-poorly organized subpopulations of Line 3) were integrated using canonical correlation analysis (CCA) in Seurat v4.0.3 (29). Only cells with between 2,000 and 6,000 unique genes (features) were included in the analysis. Only cells with < 10% of reads mapping to mtDNA-encoded genes were included. Counts data were normalized using the NormalizeData function (Seurat) with the following parameters: normalization.method= “LogNormalize”, scale.factor = 10000. 2,000 variable features were identified with the FindVariableFeatures function using the vst selection method. Integration anchors were identified using the FindIntegrationAnchors function using 25 dimensions. An assay “Integrated” was generated for the 2000 variable features using the IntegrateData function using 25 dimensions.

### Dimensionality reduction with UMAP and cell type annotation

30 principal components were identified out of the integrated dataset described above using the RunPCA function (Seurat) (29). Uniform manifold approximation and projection (UMAP) was performed using the RunUMAP function using 25 principal components. Cells were then clustered using Seurat’s FindNeighbors and FindClusters functions with the following parameters: k.param = 20, reduction=”pca”, dims=1:25 (for FindNeighbors); resolution=0.5 (FindClusters) using the original Louvain algorithm as implemented by Seurat. As shown in

**Supplemental Figure 1**, the expression patterns of genes previously shown to be markers of retinal organoid cell types were then examined in these clusters and were used to manually annotate each cluster with a cell type (30). In all calculations of photoreceptor purity, cells contained in clusters which highly express CRX were counted as fate-committed photoreceptors and all other clusters were treated as non-photoreceptor cells. Dot plot of cluster expression was generated using scCustomize (31).

**Figure 1.**
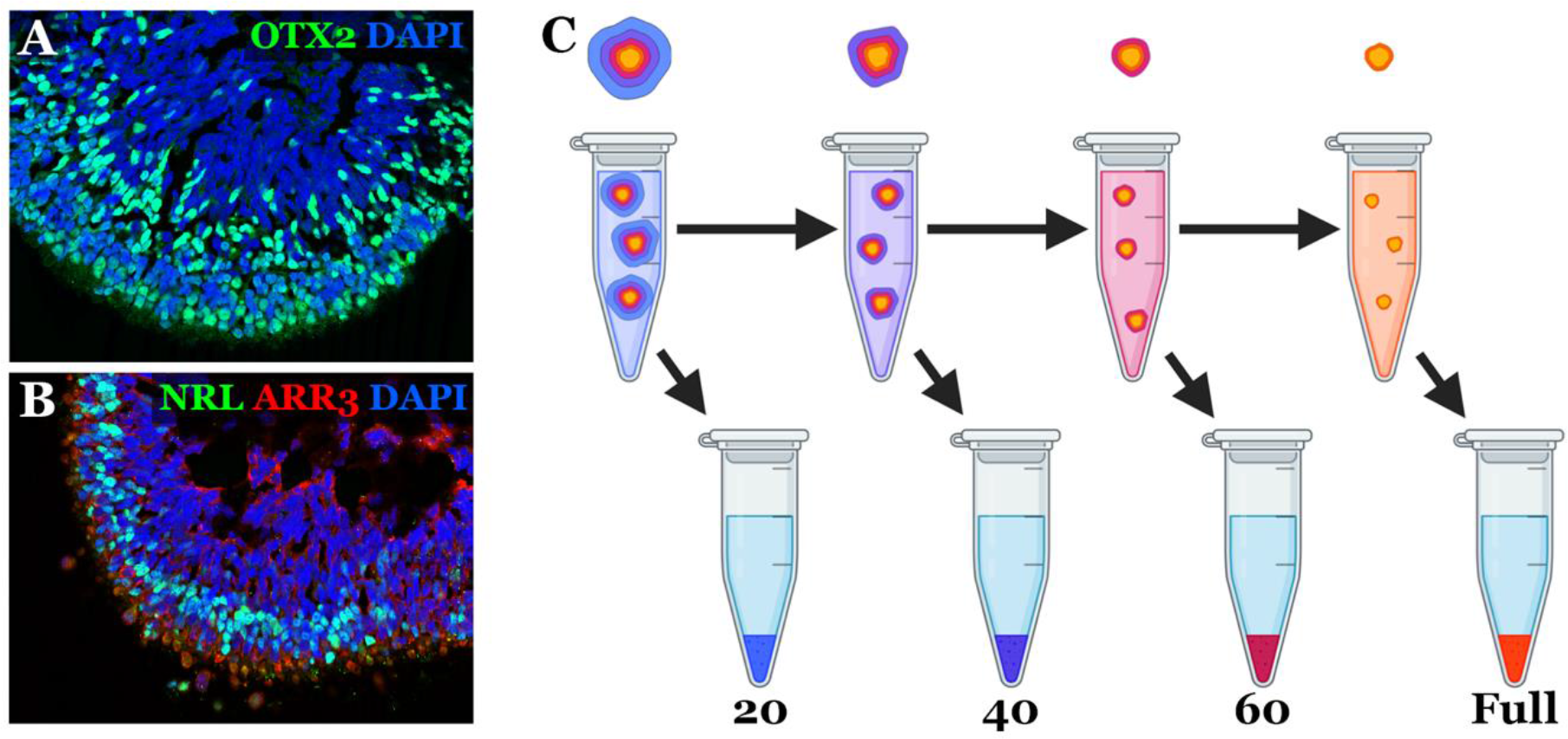
Overview. A-B) Patient iPSC-derived retinal organoids (D160) showing characteristic laminar structure with photoreceptors expressing OTX2, NRL, and ARR3 present on their outer layer. C) Partial dissociations were performed by placing organoids in papain on a rotating rack at 37°C for 20-minute increments. In between each stage of dissociation, the supernatant containing liberated cells was removed, after which the organoids were rinsed and placed in fresh papain. Following 60 minutes of total dissociation, remaining aggregates were fully dissociated via trituration.

### Immunocytochemistry

Day 160 organoids were fixed with 4% paraformaldehyde for 30-60 minutes at room temperature and equilibrated to 15% sucrose in PBS, followed by 30% sucrose. Organoids were cryopreserved in 50:50 solution of 30% sucrose/PBS: tissue freezing medium (Electron Microscopy Sciences, Hatfield, PA) and cryosectioned (15 μm). Sections were blocked with 5% normal donkey serum, 3% bovine serum albumin (BSA), and 0.1% Triton X-100 and stained overnight with the primary antibodies listed in **Table S1**. Secondary antibodies (**Table S1**) were incubated for 1 hour and cell nuclei were counterstained using DAPI (Thermo Fisher Scientific; Cat# 62248).

### Data Visualization

Plots for **Figure 2** and **Figure 3** were created using the Julia packages CSVFiles.jl, FileIO.jl, DataFrames.jl, StatsPlots.jl, ImageIO.jl, Images.jl and Colors.jl (32–34).

**Figure 2.**
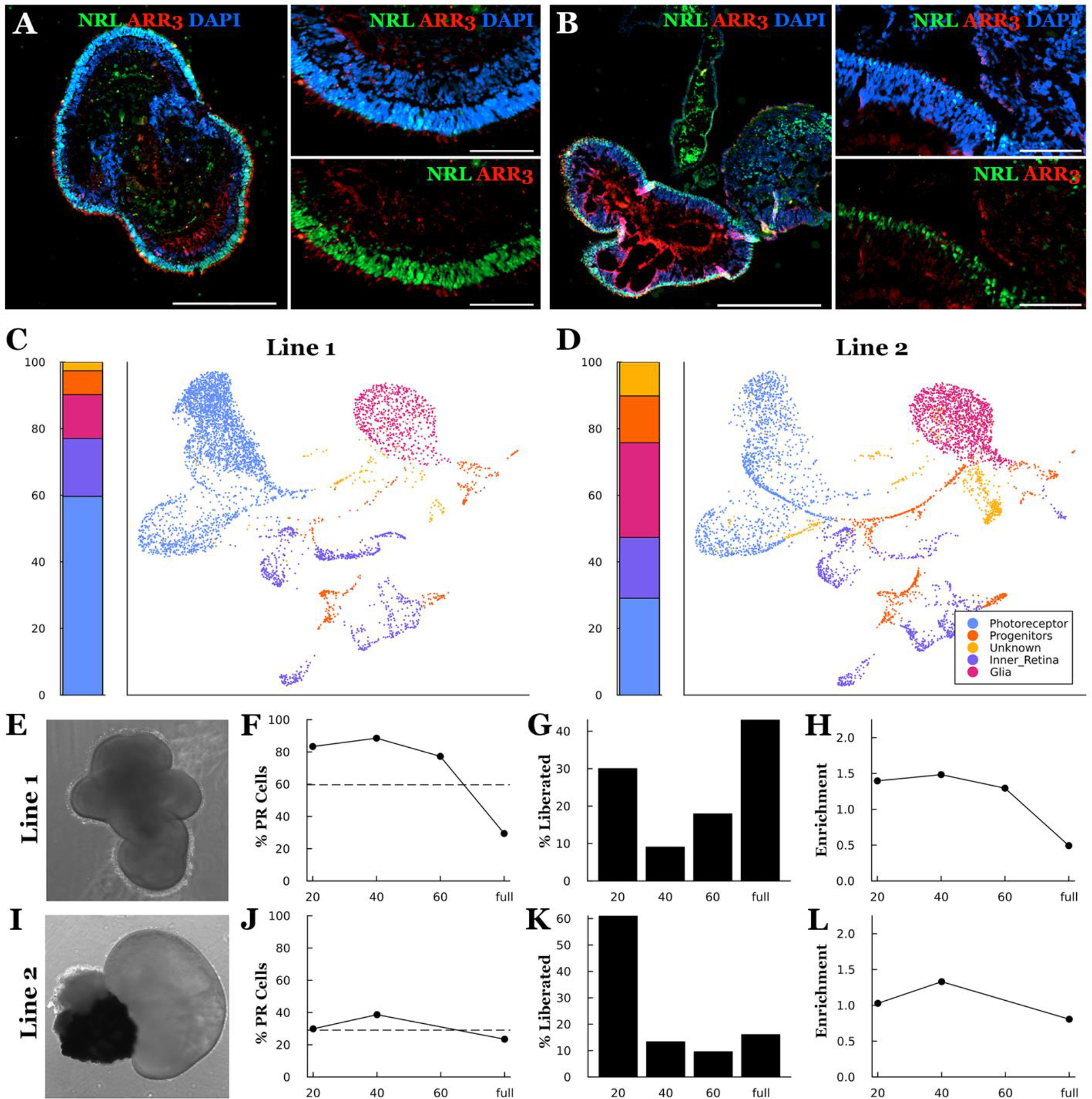
Partial dissociation of two patient lines. A-B) Characteristic structure of A) well organized Line 1 organoids and B) poorly organized Line 2 organoids (scale bar of primary panel = 500μm, insets = 100μm). C-D) Bar and UMAP plots showing the relative proportion and gene expression patterns of cells collected during control dissociations of each cell line. E-L) Photoreceptor purity and dissociation rate as a function of dissociation time. Each row shows a representative organoid (E, I) along with the photoreceptor purity (F, J) (dashed line shows the photoreceptor purity of a control dissociation of the same cell line), cell recovery (G, K), and photoreceptor enrichment (H, L) (fold change relative to a control dissociation) for each fraction recovered during staged partial dissociation. Cell fractions were taken after 20, 40 and 60 minutes of total dissociation time, after which the remaining aggregates were fully dissociated via trituration (full) and collected. The 60-minute timepoint for the purity and enrichment plots of panel (J, L) are missing due to the library preparation failing for that sample.

**Figure 3.**
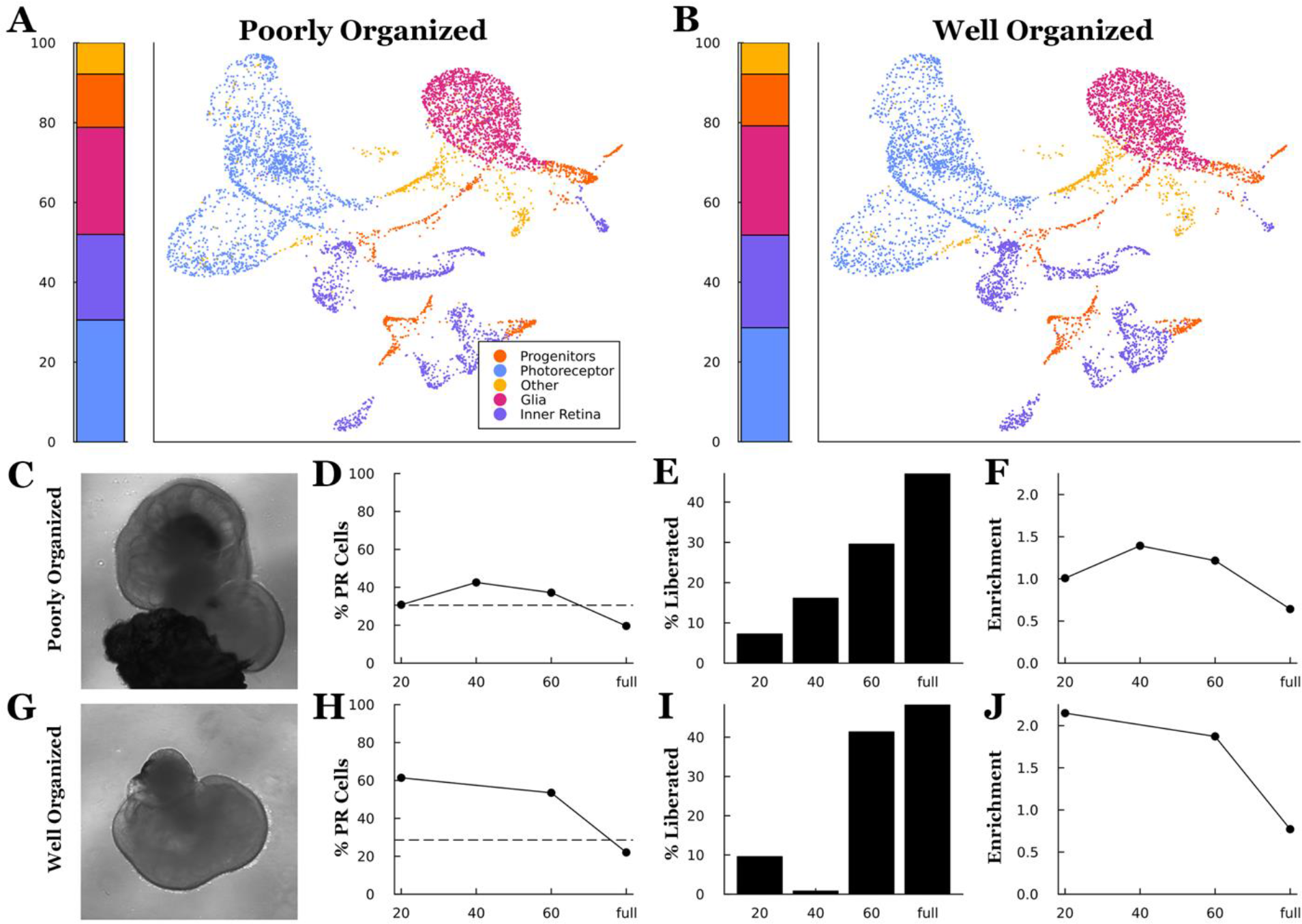
Partial dissociation of a third patient line pre-sorted by organoid organization. A-B) Prior to performing staged partial dissociation organoids produced during a single round of differentiation were sorted into ‘well organized’ and ‘poorly organized’ group Bar and UMAP plots showing the relative proportion and gene expression patterns of cells collected during control dissociations of each group of organoids. C-J) Photoreceptor purity and dissociation rate as a function of dissociation time. Each row shows a representative organoid (C, C) along with the photoreceptor purity (D, H) (dashed line shows the photoreceptor purity of a control dissociation of the same cell line), cell recovery (E, I), and photoreceptor enrichment (F, J) (fold change relative to a control dissociation) for each fraction recovered during staged partial dissociation. Cell fractions were taken after 20, 40 and 60 minutes of total dissociation time, after which the remaining aggregates were fully dissociated via trituration (full) and collected. The 40-minute timepoint is missing from the purity and enrichment panels (H, J) due to not enough cells being collected for scRNAseq analysis.

### Study approval

This study was approved by the institutional review boards of the University of Iowa and adhered to the tenets set forth in the Declaration of Helsinki. Written informed consent was obtained from all subjects prior to participation.

## Results

As shown in **Figure 1A-B**, retinal organoids produced using modern 3D differentiation protocols exhibit a characteristic laminar structure closely recapitulating normal retinal architecture. To determine if partial organoid dissociation can be used to selectively harvest photoreceptor cells, retinal organoids were incubated in papain and liberated cells were collected at the given timepoints for scRNAseq analysis. At 20-minute intervals, retinal organoids were allowed to settle, supernatant was collected, and the number of liberated cells were determined using a Moxi GO II cell counter (**Figure 1C**). Remaining organoids were rinsed and placed in fresh papain for continued dissociation. After one hour of total dissociation time, the remaining solid organoids were mechanically dissociated via trituration with a 1mL pipette tip. Retinal organoids incubated in papain for 60 minutes followed by mechanical dissociation were included as complete dissociation controls (28). Following collection of each dissociation fraction, cells were pelleted and prepared for scRNAseq. The sequenced cells were then clustered, and each cluster was annotated with a cell type corresponding to the expression pattern of the member cells. The percentage of cells present in CRX positive photoreceptor clusters was used to assess photoreceptor purity in each fraction (26, 28).

For autologous photoreceptor cell replacement, iPSC-derived retinal organoids will need to be generated on a per patient basis. While recent work has demonstrated the feasibility of this approach, variability between patient-derived iPSC lines can make clinical manufacturing difficult. Specifically, not all patient derived iPSC lines are able to generate retinal organoids of the same quality or number (35). For instance, Line 1 is an efficient producer of laminated retinal organoids that contain many photoreceptor cells within their outermost layer (**Figure 2A**). Line 2 however, while still an efficient retinal organoid producer, generally gives rise to organoids that are less well organized, and often contain large aggregates of retinal pigmented epithelium (RPE) (**Figure 2B)**. Despite these structural differences, both lines contained the same collection of cell types with similar gene expression patterns (**Figure 2C-D**). After subjecting both lines to staged partial dissociation, we found that the well-organized Line 1 yielded more cells in enriched fractions (i.e. fractions with photoreceptor cell purity greater than control dissociations) than the Line 2 organoids (**Figure 2E-F**). Importantly, relatively large numbers of cells were recovered in these enriched fractions, indicating that this enrichment strategy is capable of isolating photoreceptor cells at reasonable total yield. While a substantial number of cells were recovered from the enriched fractions of both lines, there is a difference between their dissociation kinetics. In the well-organized line, a large number of photoreceptor cells were liberated in the first fraction, after which the dissociation rate slowed before eventually increasing again (**Figure 2E**). One possible explanation for this behavior is that dissociation stalled when the outer plexiform layer (OPL) beneath the outer layer of photoreceptor cells was reached. We hypothesize that the less organized Line 2 does not exhibit the same dissociation kinetics because the attached clusters of RPE and general disorganization impacts the integrity of the OPL (e.g., once the RPE cluster is removed the inner layers of the organoid are exposed allowing non-photoreceptor cells to escape).

To confirm that the differences in maximum purity and enrichment that we observed between the Line 1 and Line 2 were due to organoid structure, we proceeded to perform partial dissociations on organoids generated from a third line which has both well and poorly organized retinal organoids. While both the well-organized (**Figure 3A**) and poorly organized (**Figure 3B**) populations contained the same relative proportions of the constituent cell types, the well-organized organoids yielded dissociation fractions that were more highly enriched for photoreceptor cells (**Figure 3C-D**). Further, the well-organized sample showed the same characteristic dissociation kinetics as Line 1, indicating that this signature is likely a result of the highly organized structure of the organoids as hypothesized above. While less efficient in terms of total yield, these results also indicate that our partial dissociation approach could be used to produce highly enriched photoreceptor cell fractions from patient lines that produce organoids with varying levels of organization by mechanically pre-sorting organoids prior to dissociation.

## Discussion

Photoreceptor replacement therapies show great promise for restoring vision in patients with many forms of currently untreatable retinal degenerative blindness. cGMP-compliant 3D differentiation protocols that can produce retinal organoids from patient-derived iPSCs have been developed (9–19). In these 3D differentiation approaches, the development of retinal organoids mimics that of the human retina in utero, resulting in laminated tissue containing both committed photoreceptor lineages, which are desired for transplant, along with other retinal cell types. Given that only photoreceptors are expected to contribute to the therapeutic outcome of photoreceptor replacement therapies, techniques capable of enriching photoreceptors from dissociated retinal organoids are needed to increase the potency of these treatments. In this work, we have shown that partial dissociation can be used to exploit the structural organization of retinal organoids to produce highly pure populations of dissociated photoreceptor cells without the use of specialized equipment or antibody-based labels. While our initial results have been encouraging, further development is needed to ensure that this approach produces consistent results across lines derived from the many patients in need of photoreceptor cell transplants. The most obvious limitation of our partial dissociation technique is that its performance is dependent on the degree of organization present in the organoids of interest. While we have shown that this issue can be overcome by simply preselecting organoids with obvious lamination and absence of disorganized appendages, the level of waste resulting from this strategy is not desirable. This is especially true as large numbers of differentiations need to be performed in parallel to produce autologous therapies for many patients in a clinical setting. Therefore, we and others have been investigating protocol changes aimed at producing highly organized retinal organoids. For instance, by using 96 well u-bottom plates, Harkin et al. recently demonstrated a highly effective iPSC reaggregation strategy that enabled production of high quality uniform embryoid bodies. By plating these embryoid bodies on an adherent 2D surface and allowing for a short period of expansion and retinal cell fate commitment, the authors were able to pick optic vesicles, and generate laminated retinal organoids with a uniform shape and size from all iPSC lines evaluated (36). At later stages of development retinal organoid aggregation and death of inner retinal neurons can result in structural changes that may not be conducive to the partial dissociation strategy presented in this study. The death of inner retinal neurons can be attributed to a lack of nutrient and gas exchange as large organoids sit in static culture conditions, which also allow organoids to aggregate. To address these concerns, we recently began to explore the use of bioreactor systems similar to those that have been previously reported (37–39). Clinostat bioreactors such as the ClinoStar (CelVivo) use slow rotation to eliminate settling, increase perfusion, and reduce organoid aggregation, enabling accelerated production of large, laminated organoids, that may be ideally suited to the partial dissociation strategy described in this study (**Figure 4**).

**Figure 4.**
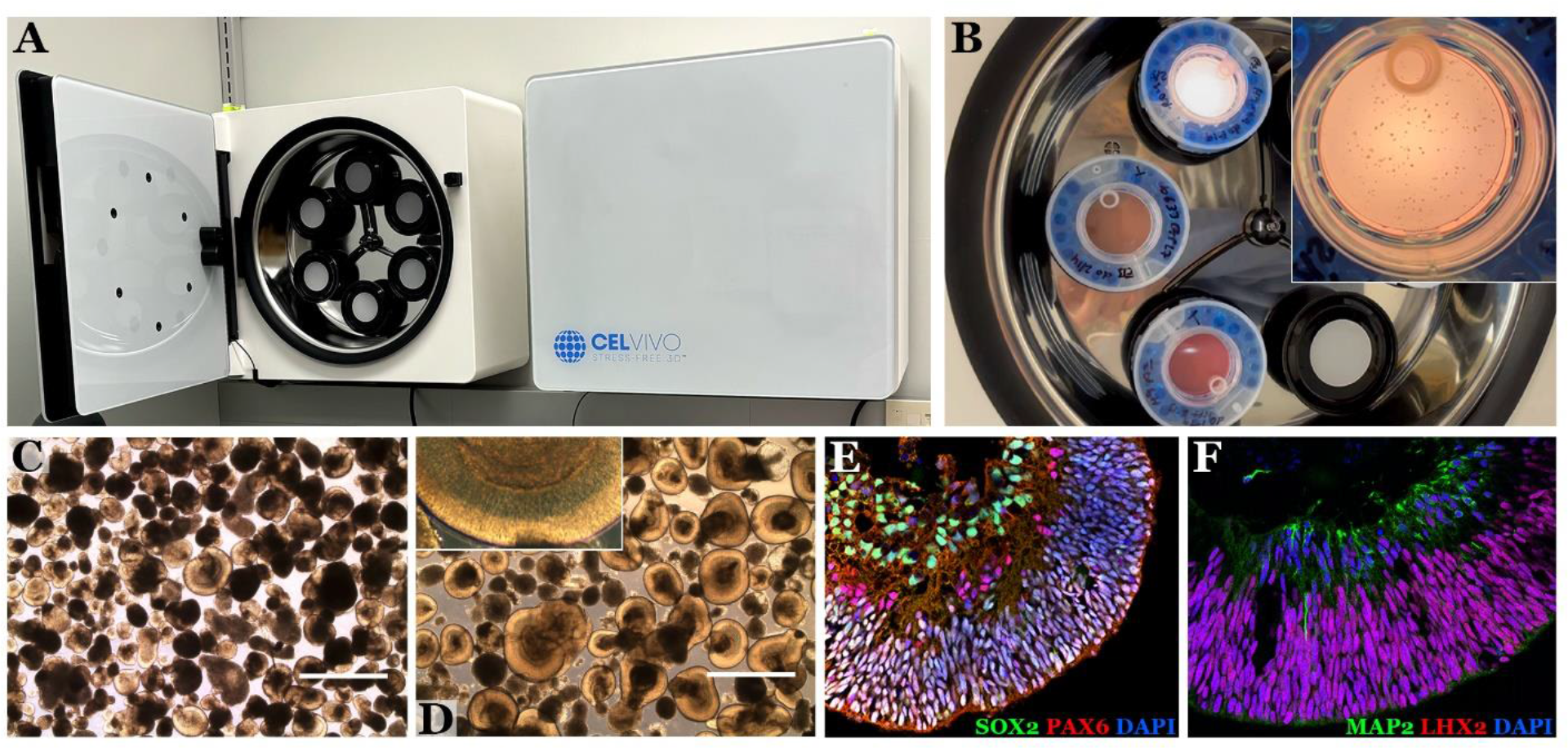
Clinostat bioreactors produce highly organized retinal organoids. A-B) Clinostar bioreactors simulate microgravity by continuously rotating several individual culture vessels. This continuous rotation appears to increase organoid perfusion and reduce aggregation, resulting in D-F) larger, better organized and more mature organoids as compared to C) traditional 3D differentiation techniques.

Given optimal organoid structure, one additional consideration is that organoids derived from different patients or produced during different rounds of differentiation may contain differing proportions of photoreceptors or dissociate at different rates. In this case, simply dissociating organoids for the same amount of time may not be sufficient to ensure that the composition of liberated cells is consistent across patients and manufacturing runs. However, our initial results indicate that there may be another indicator which could be used to control for batch-to-batch variability in dissociation rate. In all the dissociations that we performed on well-organized retinal organoids, dissociation rate first decreased from its initial value before increasing. This suggests that dissociation kinetics could be used to monitor the process in a time-independent manner, allowing for production of consistent cell populations across patients and rounds of differentiation.

## Conclusions

While retinal organoids are an ideal source of cells for autologous photoreceptor cell replacement, the fact that state of the art differentiation strategies result in complex tissues containing multiple cell types complicates their potential clinical use. To ensure uniform, positive treatment outcomes, it is important that consistent cell populations be used for transplant, and that these populations only contain cells expected to contribute to the recovery of vision. While the need for clinically compatible photoreceptor enrichment strategies is clear, most traditional approaches have significant drawbacks. FACS and MACS, while mature technologies, require the use of expensive equipment and the exposure of cells destined for transplant to antibody-conjugated labels. In this work, we demonstrate how partial dissociation can be used to exploit the structure of iPSC-derived retinal organoids to produce highly pure photoreceptor cells without using specialized equipment or reagents such as antibody tags. Although our current protocols do not always produce perfectly laminated organoids from every line of patient-derived iPSCs, protocol improvements including clinostat bioreactors and pre-sorting of laminated organoids from populations of mixed quality show promise for increasing the performance and applicability of this technique. In addition, we believe that monitoring dissociation kinetics will provide additional feedback to reduce line-to-line variability and provide additional documentation as to the integrity of the cell fractions liberated. In summary, partial dissociation constitutes a cheap, clinically translatable, and effective technique for purification of transplantable photoreceptor cells.

