## supplemental material for "Device-free isolation of photoreceptor cells from patient iPSC-derived retinal organoids"

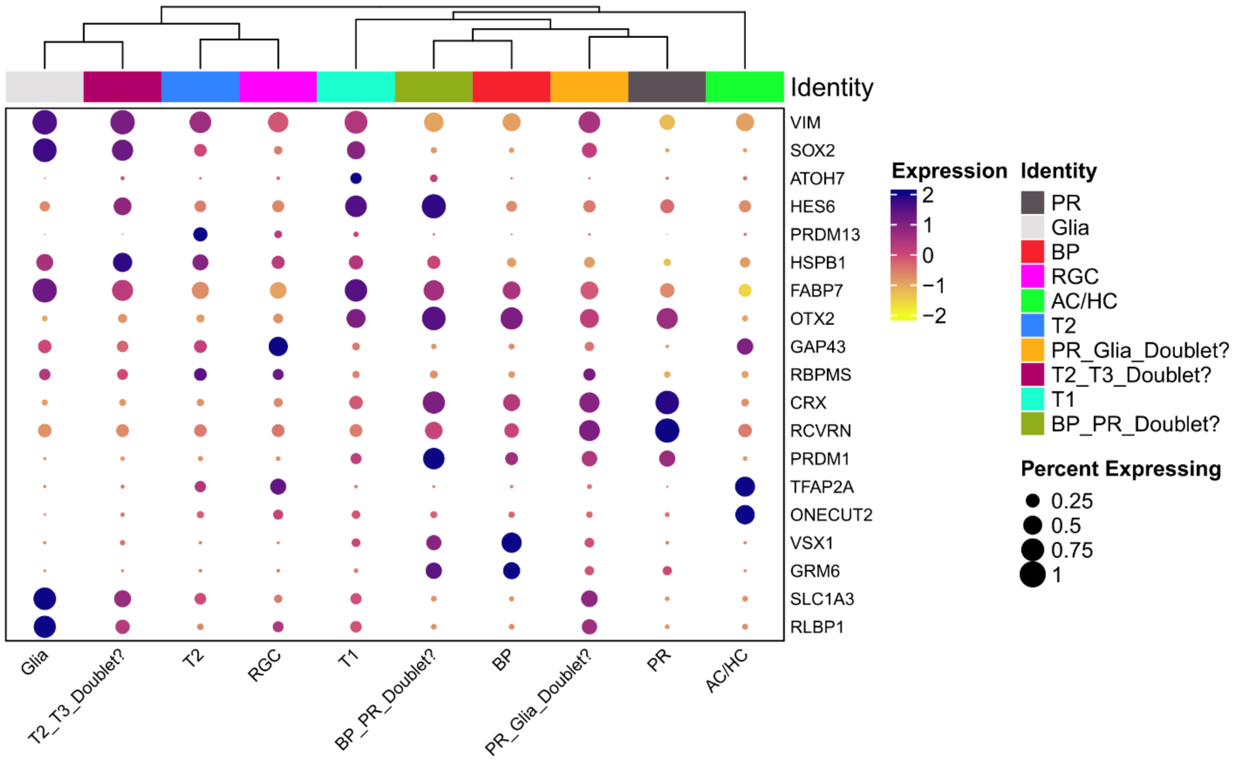


Supplemental Figure 1: Gene expression and cell type annotations for clusters identified by Seurat. Fate-committed photoreceptor clusters were identified through their expression of canonical markers such as CRX and RCVRN.

Table S1: Antibodies used

| Primary Antibodies | | | |
|---|---|---|---|
| Target | Host | Vendor | Catalog number |
| ARR3 | Rabbit | LifeSpan Bio | LS-C368677 |
| NRL | Goat | R&D Systems | AF2945 |
| OTX2 | Goat | R&D Systems | AF1979 |
| Secondary Antibodies | | | |
| Donkey anti-goat 488 | | Thermo | A11055 |
| Donkey anti-rabbit 647 | | Thermo | A31573 |
